# An AI-driven approach for nanobody affinity maturation

**DOI:** 10.1101/2025.11.11.687768

**Authors:** Xuceng Yang, Ziang Guo, Qi Zhao

## Abstract

B7-H3 (CD276), an immunoregulatory checkpoint molecule overexpressed in numerous cancers, is a promising therapeutic target. Nanobodies possess unique advantages for targeting B7-H3, such as small size, high stability, and the ability to bind cryptic epitopes. However, the rational affinity maturation of these nanobodies is challenging, especially in the absence of detailed structural data on antigen-antibody interactions. Here, we present a computational strategy that leverages artificial intelligence (AI) and molecular modeling, including homology modeling, molecular docking, and free-energy calculations—to systematically predict affinity-enhancing mutations for humanized anti-B7-H3 nanobodies. This AI-driven framework provides a powerful and cost-effective pipeline for accelerating the development of high-affinity nanobody therapeutics prior to experimental validation.

## Introduction

Nanobodies are the single-domain antigen-binding fragments derived from the heavy-chain–only antibodies of camelids. Composed of a single variable domain (VHH), nanobodies typically have a molecular weight of ∼12–15 kDa, display high solubility and thermal stability, and retain full antigen-binding capacity despite their small size[1]. These properties enable nanobodies to penetrate dense tissues, recognize cryptic or recessed epitopes, and be readily engineered for therapeutic, diagnostic, and imaging applications. Several nanobody-based drugs have entered clinical trials or received regulatory approval, underscoring their potential as next-generation antibody therapeutics.

The development of therapeutic antibodies traditionally begins with the identification of target antigens and the generation of antigen-specific binders through methods such as hybridoma technology, phage or yeast display libraries, or single B-cell cloning. Once a lead antibody is obtained, structural characterization of the antigen–antibody complex becomes crucial for guiding rational optimization. High-resolution structures are typically determined by X-ray crystallography or, more recently, single-particle cryo-electron microscopy (cryo-EM), which reveal the precise arrangement of complementarity-determining regions (CDRs) at the binding interface. Mapping of the epitope—the antigenic surface recognized by the antibody— is a critical step for understanding binding mechanisms and improving therapeutic performance. Classical approaches include hydrogen–deuterium exchange mass spectrometry (HDX-MS)[2], site-directed mutagenesis (“alanine scanning”), peptide microarrays[3], and surface plasmon resonance (SPR)[4] competition assays, all of which can reveal contact residues and energetic “hot spots” at the antibody–antigen interface. Combining epitope information with paratope analysis allows researchers to pinpoint key interactions that determine binding affinity and specificity. Following epitope identification, antibodies are typically optimized through affinity maturation. Traditional affinity maturation relies on directed evolution techniques such as error-prone PCR, DNA shuffling, or display-based selection to introduce and screen large numbers of random mutations in the CDRs, selecting variants with improved binding.

In cases where experimental structures are unavailable, homology modeling, AlphaFold predictions, or integrative molecular modeling can provide reliable three-dimensional representations of both antigens and antibody variable domains. More recent strategies incorporate rational design, in which specific residues are mutated based on structural insights, or computational protein design, which predicts mutations predicted to lower binding free energy. These approaches are often complemented by molecular dynamics (MD) simulations[5], which capture the conformational flexibility of antibodies and antigens and provide a more accurate assessment of energetic effects. Despite these advances, conventional antibody optimization remains resource- and time-intensive, requiring extensive wet-lab screening to identify beneficial mutations among a vast combinatorial space. The increasing accuracy of in silico methods—including molecular docking, free-energy calculations, and machine-learning– based predictors—offers a cost-effective alternative to accelerate antibody engineering, reduce experimental workload, and enable the rational design of variants with enhanced affinity or altered specificity. Molecular dynamics (MD) simulations and related computational approaches have become indispensable tools for predicting antibody–antigen interactions, estimating binding free energies, and guiding rational mutagenesis.

However, significant limitations remain that hinder their broad adoption in therapeutic antibody development. First, the field is fragmented by a large and diverse set of software platforms—including commercial packages and open-source tools—each with different force fields, scoring functions, and parameterization schemes. These variations often lead to inconsistent results across studies, making it difficult to reproduce or directly compare predictions. Second, there is no universally accepted workflow for integrating homology modeling, docking, alanine scanning, saturation mutagenesis, and stability assessment. Researchers frequently combine disparate tools with customized scripts, resulting in ad hoc protocols that require substantial expertise and are challenging for experimental laboratories to implement.

To address these gaps, we established a standardized, end-to-end computational pipeline that integrates structure modeling, docking refinement, energy-based mutational scanning, and stability filtering within a single coherent framework. This protocol provides clear decision criteria at each stage, facilitating reproducibility and practical application. As a proof of concept, we applied the workflow to the rational engineering of anti–B7-H3 single-domain antibodies (VHHs) and experimentally validated the predicted mutations, demonstrating both the feasibility and predictive power of this streamlined approach.

B7-H3 (CD276) is a type I transmembrane glycoprotein belonging to the B7 family of immune checkpoint molecules[6]. It is broadly overexpressed on a variety of human tumors— including breast, lung, kidney, and neuroblastoma—while exhibiting limited distribution in normal tissues[7], [8]. Recent studies have revealed that, in addition to its role in tumor immune evasion, angiogenesis, and metastasis, B7-H3 can directly interact with the oncogenic receptor tyrosine kinase c-Met, activating the c-Met/STAT3 signaling pathway to promote cancer cell stemness. These findings uncover a tumor cell–intrinsic mechanism and provide new therapeutic opportunities for B7-H3–targeted immunotherapy[9]. Specifically, we first humanized the nanobody G1, which recognizes the tumor-associated antigen B7-H3. Upon humanization, the binding affinity of G1E1 decreased markedly, prompting us to restore and enhance its affinity through in silico identification of beneficial mutations. The predicted variants were subsequently validated by ELISA and BLI assays, confirming the reliability of the computational predictions. Notably, B7-H3 was deliberately chosen as the model system to rigorously test the robustness of our *in silico* workflow, as no experimentally determined crystal structure is available for this antigen. Despite this limitation, our computationally guided design successfully achieved a measurable improvement in antibody affinity, underscoring the practical applicability and reliability of the workflow even in the absence of high-resolution structural data.

## Method

### Generation of Antibody and antigen structure model

Because no experimentally determined crystal structure of human B7-H3 is currently available, we constructed structural models of the antigen and nanobodies using a combination of homology modeling and AI-based prediction. The murine B7-H3 ectodomain (PDB ID: 4I0K)[10], which adopts a 2-Ig–like architecture, was first retrieved from the Protein Data Bank and employed as a template in SWISS-MODEL[11], [12] to generate a homology model of human B7-H3. Given that the predominant human isoform is a 4-Ig–like form, and that the R1 and R2 Ig domains share high sequence similarity, this murine template provided a suitable starting point for modeling the R2-containing regions of the human protein. In parallel, an independent full-length human B7-H3 structure was predicted using AlphaFold[13], and the resulting model was subsequently re-refined within SWISS-MODEL to improve local geometry and stereochemical quality. The single-domain antibodies (nanobodies) and their humanized versions G1E1 were modeled in SWISS-MODEL using high-resolution VHH structures from the PDB as templates.

### Structure evaluation and refinement

During structural evaluation and refinement, the models were first assessed using three key indices automatically calculated by SWISS-MODEL during modeling. These parameters provide an initial quality estimation before further validation. The Global Model Quality Estimate (GMQE) combines target–template alignment features and template structure properties through a neural network to predict the local distance difference test (lDDT) score of the resulting model. GMQE offers a preliminary assessment before model building and is later updated after model generation by incorporating the QMEANDisCo global score for improved reliability. The QMEANDisCo global score reflects the average per-residue agreement between the model and evolutionary/structural features and correlates well with the lDDT score; higher values indicate better overall model accuracy[14].

To further validate and refine the modeled structures, the final PDB files were uploaded to SAVES v6.1 (https://saves.mbi.ucla.edu/), which integrates additional quality assessment tools including ERRAT[15], VERIFY 3D[16], [17], and the Ramachandran (Rama) plot[18]. ERRAT analyzes non-bonded atom–atom interactions and reports an overall quality factor[15], where values above 90% indicate a reliable structure and errors are ideally minimized in critical regions such as the nanobody CDR loops. VERIFY 3D evaluates the compatibility of each amino acid residue with its 3D environment[15]; a model is generally considered high quality when more than 80% of residues achieve a score above 0.2. The Rama plot examines backbone dihedral angles (φ and ψ) to ensure residues occupy favorable conformational space, with a high percentage of residues in favored regions reflecting accurate stereochemistry. If any model failed to meet these quality thresholds, the structure was iteratively refined by re-submitting it to SWISS-MODEL for template-guided rebuilding or by performing local side-chain refinement in Discovery Studio, particularly targeting residues within the CDRs. Models that consistently failed to improve beyond acceptable scoring criteria were rebuilt using alternative templates to ensure optimal structural reliability.

### Local docking of Antigen-antibody complex

The docking calculations were performed in Discovery Studio (DS)[19]using the *Dock Protein* module, which provides the rigid-body docking algorithm ZDOCK and the refinement docking algorithm RDOCK. ZDOCK, developed at the University of Massachusetts Medical School, is a rigid-body protein–protein docking algorithm based on a fast Fourier transform (FFT) correlation approach. This method efficiently searches the translational and rotational space of two proteins to predict possible binding orientations. During docking, ZDOCK generates approximately 2,000 candidate poses, which are automatically grouped into about 200 clusters plus additional unclustered individual poses. Each cluster is ranked by size, and the docking results are scored using the ZDOCK scoring function. The ZDOCK score reflects shape complementarity between the two proteins, with higher scores indicating more favorable fits. ZDOCK is also available as a free web service[20] (https://zdock.wenglab.org/). From the 2,000 initial poses, the top poses within the largest clusters were examined, and typically the best ∼20 poses were selected for further optimization (the exact number can be adjusted according to the need). These selected poses were processed using the *Process Poses (ZDOCK)* module to remove steric clashes and prepare structures for refinement.

Refinement and rescoring were carried out using RDOCK, which applies a CHARMm polar H based energy minimization protocol to improve the near-native quality of docking predictions. RDOCK performs a two-stage energy minimization, first removing steric clashes and then evaluating electrostatic and desolvation energies. The final ranking of poses is based on the sum of these energy terms, allowing discrimination of the most stable binding conformations. This refinement helps correct for limitations of the rigid-body ZDOCK method by optimizing side-chain packing and electrostatic interactions.

Before RDOCK refinement, the resulting complexes of ZDock were carefully inspected and filtered. Typically, top-ranked poses from the first five clusters were retained to ensure structural diversity, and occasionally unclustered high-scoring poses (cluster 2001) were included if they showed unique binding orientations. The final set of refined complexes was used as the predicted antigen–antibody docking models.

### Alanine Scanning and Single-Mutation Analysis *in silico*

Protein mutational “hotspots” are often identified through alanine scanning, a strategy in which specific amino acid residues are individually substituted with alanine to evaluate their contribution to protein function or binding[21]. Alanine is ideal for this purpose because of its small side chain (a simple methyl group), chemical inertness, and minimal effect on the protein’s secondary structure. By systematically replacing residues with alanine, it is possible to determine whether a particular side chain plays a critical role in maintaining the biological activity or binding affinity of the protein. Although alanine scanning is traditionally performed experimentally through site-directed mutagenesis or PCR-based mutant libraries, here the process was carried out *in silico*.

For nanobodies, which have relatively small sequences, all residues can be subjected to alanine scanning. For larger antibodies (e.g., IgG) or antibodies with poorly defined paratopes, computational cost may become limiting, and scanning is typically focused on the complementarity-determining regions (CDRs) or on residues located at the antigen–antibody interaction interface, as these residues are most likely to directly influence antigen recognition while maintaining the structural core of the antibody. Each mutation is evaluated by calculating the change in binding free energy, expressed as ΔΔG, which represents the difference in predicted binding free energy between the mutant and wild-type complexes (ΔΔG = ΔG_mutant – ΔG_wildtype)[22]. A negative ΔΔG indicates that the alanine mutation enhances binding (i.e., lowers binding free energy), whereas a positive value suggests a loss of binding affinity.

It is noteworthy that two complementary strategies are commonly used to interpret alanine-scan results for guiding mutational design. In the first strategy, residues whose substitution to alanine produces a positive ΔΔG (i.e., mutation destabilizes the complex and reduces affinity) are designated as binding hotspots; such residues make large favorable energetic contributions to the interaction, and targeted replacement of these positions with alternative side chains that form stronger interactions can yield substantial affinity gains. The second strategy focuses on residues for which alanine substitution yields a negative ΔΔG (i.e., the alanine mutant shows increased binding affinity)[23]. These positions indicate that the wild-type side chain may exert a small unfavorable effect on binding, so replacing that residue with an inert side chain (such as alanine) or other benign substitutions can modestly improve affinity without large structural perturbation. Although changes identified by this second approach typically confer smaller improvements than mutations targeting classical hotspots, they carry lower risk of disrupting folding or stability and therefore represent conservative, low-risk optimization sites; strictly speaking, ΔΔG < 0 sites are not canonical “hotspots” but rather residues with negative contributions that are amenable to optimization.

Based on the alanine scanning results, the top candidate residues with ΔΔG < 0 were selected— typically the ten most favorable sites—for further saturation mutagenesis. At each hotspot, all 19 alternative amino acids were computationally substituted, and the resulting mutants were again evaluated for ΔΔG. Mutations yielding negative ΔΔG values were identified as promising candidates for improving antigen binding, as they indicate side chains that may provide more favorable steric, electrostatic, or conformational interactions than alanine or the original residue.

### Stability Calculation and Multipoint Mutation

Following the initial alanine scanning and single-site saturation mutagenesis, candidate mutations with ΔΔG < –1.0 kcal/mol (indicating a strong predicted increase in antigen-binding affinity) were selected for stability analysis. The structural stability of each mutant was evaluated in Discovery Studio using the “Calculate Mutation Energy (Stability)” module, which estimates the free energy change associated with the mutation (ΔΔG_stable). A negative ΔΔG_stable value suggests that the mutation enhances the thermodynamic stability of the antibody, whereas a positive value indicates a potential destabilizing effect. Only mutations predicted to both increase binding affinity (ΔΔG < –1.0 kcal/mol) and improve stability (ΔΔG_stable < 0) were retained as promising candidates.

The top-ranked single-site mutations were then combined to explore potential multipoint mutants, as synergistic effects among multiple favorable substitutions can yield greater improvements in binding affinity than individual mutations alone. Discovery Studio provides built-in tools to model double and triple mutations, allowing systematic exploration of two or three-residue combinations. For each multi-mutation construct, both binding affinity (ΔΔG) and structural stability (ΔΔG_stable) were recalculated to identify combinations that maintained or improved antibody stability while further enhancing antigen binding. This approach enables the rational design of multi-site variants with optimized affinity–stability profiles, while avoiding destabilizing mutations that might compromise expression or folding in subsequent experimental validation.

### Integration of Multiple Docking Conformations and Re-mutation Analysis

Because RDOCK typically generates several energetically favorable docking conformations (poses), we focused on complexes derived from the top-ranking clusters for virtual mutagenesis. Each selected pose can exhibit subtle differences at the antigen–antibody interface, which may lead to distinct sets of favorable mutations. To ensure that the final mutation strategy is not biased by a single docking solution, beneficial mutations were compared across all selected complexes. Mutations or residue positions that repeatedly emerged as advantageous in multiple independent poses were considered high-confidence candidates, as their recurrence suggests a consistent energetic contribution to antigen binding regardless of small conformational differences. High-frequency mutations identified from this cross-pose comparison were then introduced into all selected docking complexes to further validate their effects. For each mutant, changes in binding affinity (ΔΔG) and structural stability (ΔΔG_stable) were calculated within each pose, and the average values across poses were determined. This averaging process helps mitigate pose-specific artifacts and highlights mutations that provide robust improvements in both affinity and stability across different plausible antigen–antibody geometries. Finally, combinations of mutations with the lowest mean ΔΔG values and increased stability profiles were prioritized as the most reliable candidates for experimental validation.

### Expression of nanobodies

Based on the computational analysis, two double-point mutants were selected. To enable downstream expression and functional testing, the corresponding DNA sequences were synthesized commercially. Each nanobody coding sequence was flanked with SfiI restriction sites to facilitate directional cloning. The synthesis company inserted the gene fragments into a pUC-GW-Amp cloning vector and delivered the recombinant plasmids for subsequent subcloning.

The recombinant plasmids were transformed into *E. coli* TOP10 competent cells for amplification, and the plasmid DNA was subsequently purified using a commercial plasmid extraction kit. The nanobody genes were released from pUC-GW-Amp by SfiI digestion and ligated using T4 DNA ligase into a pre-linearized pComb3X phagemid vector. pComb3X is widely used for phage display of single-domain antibodies and contains a lac promoter and lac operator that allow IPTG-inducible transcription. The construct also includes an OmpA signal peptide, which directs the expressed nanobody to the periplasmic space of *E. coli* to promote correct folding and disulfide-bond formation. Downstream of the nanobody coding region, a 6×His tag and a FLAG epitope were incorporated to facilitate purification and detection by affinity chromatography or immunoblotting. This cloning strategy ensures efficient production of each nanobody mutant and provides versatile tags for both biochemical characterization and subsequent binding assays.

The recombinant pComb3X plasmids encoding the nanobody variants were transformed into *Escherichia coli* HB2151 cells for periplasmic expression. Protein expression was induced with isopropyl β-D-1-thiogalactopyranoside (IPTG) when cultures reached the appropriate optical density. After induction, the bacterial pellets were harvested and treated with polymyxin B to selectively disrupt the outer membrane and release the periplasmic contents. The nanobody proteins were purified from the supernatant using Ni–NTA affinity chromatography, which captures the C-terminal His tag. Elution was performed with imidazole-containing buffer, and the eluates were subsequently concentrated and buffer-exchanged into phosphate-buffered saline (PBS) using ultrafiltration. Protein size and purity were verified by SDS–PAGE and NATIVE-PAGE analysis.

### Protein Purification and Purity Assessment by Size Exclusion Chromatography (SEC)

To further purify the recombinant protein and assess its homogeneity, size exclusion chromatography (SEC) was performed using an ÄKTA FPLC system equipped with a Superdex® 200 Increase 10/300 GL column (Cytiva). The column was pre-equilibrated with phosphate-buffered saline (PBS, pH 7.4) before sample loading. The protein sample was injected onto the column and eluted isocratically at a constant flow rate while monitoring absorbance at 280 nm. The elution profile was monitored to distinguish the main target protein peak from minor impurity peaks. Fractions corresponding to the principal elution peak were collected for further analysis. To assess the effectiveness of the purification, both the initial protein sample and the SEC-purified fractions were analyzed by Native-PAGE under non-denaturing conditions. The improvement in purity was evaluated by comparing the clarity and intensity of the target protein band between the original sample and the SEC-purified fractions, confirming the removal of contaminating species and the integrity of the purified protein.

### Enzyme-linked immunosorbent assay (ELISA)

The binding activity of each nanobody variant toward the extracellular domain of human B7-H3 was quantified using an enzyme-linked immunosorbent assay (ELISA). 96–well plates were coated with 50 ng/well of B7-H3 and incubated overnight at 4 °C. Plates were blocked with 3% (w/v) skim milk in PBS for 30 minutes at 37℃ to reduce nonspecific binding. Nanobody samples—including wild-type G1-cam, humanized G1E1, and 2 computationally designed mutants G1, G2—were serially diluted in blocking buffer to generate eight concentration gradients ranging from 1000 nM to 10⁻⁴ nM and incubated on the coated plates for 30 minutes at 37℃. After washing four times with 0.05% PBST (PBS with 0.05% Tween-20), an anti-FLAG monoclonal antibody was added as the detection reagent and incubated for 30 minutes at 37℃. Plates were washed again four times with 0.05% PBST, and binding was detected using 3,3′,5,5′-tetramethylbenzidine (TMB) substrate. The enzymatic reaction was stopped after approximately 2 min with 2M H_2_SO_4_ solution, and absorbance was recorded at 450 nm using a microplate reader. All measurements were performed in duplicate to ensure reproducibility. Following the initial analysis, a second independent ELISA was carried out under identical conditions to validate and directly compare the binding affinities of the nanobody variants toward B7-H3.

### BLI (Biolayer Interferometry) Analysis

Biotinylation of the protein was performed at a molar ratio of 3:1 (biotin:protein) using an NHS-ester-based biotinylation reagent. The reaction was carried out for 30 minutes at room temperature, followed by quenching with excess reagent. The biotinylated protein was then purified using a Ultrafiltration Spin Columns to remove unreacted biotin and buffer components. For biolayer interferometry (BLI) analysis, the biotinylated nanobody was immobilized onto a streptavidin (SA) biosensor (FortéBio). The sensor was first equilibrated in assay buffer containing 0.1% BSA and 0.02% PBST (PBS with 0.02% Tween-20) to establish Baseline 1. The sensor was then loaded with 200 µL of biotinylated antibody at a concentration of 2.6 µg/mL, followed by a second equilibration step (Baseline 2). The association phase was performed by exposing the loaded sensor to antigen (B7-H3) solutions at varying concentrations (100 nM, 75 nM, 50 nM, 37.5 nM, 18.8 nM, 9.8 nM, 5 nM and 0 nM) for 180 seconds. Dissociation was subsequently monitored by transferring the sensor into assay buffer. The association and dissociation curves obtained were fitted using a 1:1 binding model to determine the association rate constant (ka), dissociation rate constant (kd), and equilibrium dissociation constant (KD).

### Circular Dichroism (CD) Spectroscopy

Circular dichroism (CD) spectroscopy was employed to evaluate the secondary structure and thermal stability of the purified proteins. Protein samples were diluted to a final concentration of 0.3 mg/mL in phosphate-buffered saline (PBS, pH 7.4) and analyzed using a Chirascan CD spectrometer (Applied Photophysics, UK). Spectra were recorded from 190 to 260 nm with a 1 nm bandwidth and 1 nm data interval quartz cuvette while the temperature was increased from 20 °C to 100 °C at a heating rate of 2 °C per minute under a continuous nitrogen purge. The resulting melting curves were fitted using the Global3 analysis software (Applied Photophysics) to determine the melting temperature (Tₘ), defined as the midpoint of the unfolding transition.

## Result

### Workflow of Protocol

The overall workflow of this computational protocol is illustrated in the accompanying Figure 1. In the first step, the user provides only the amino-acid sequences of the antibody and the target antigen. The SWISS-MODEL server automatically searches the PDB and AlphaFold databases for homologous structures and presents candidate templates together with sequence-identity scores, GMQE values, and other relevant metrics. After inspecting these results, the user can select one or more suitable templates and initiate homology modeling. When the sequence identity between the query and template exceeds approximately 40%, the proteins are likely to belong to the same structural family, and homology modeling is expected to yield a reliable three-dimensional model[24].

**Figure 1:**
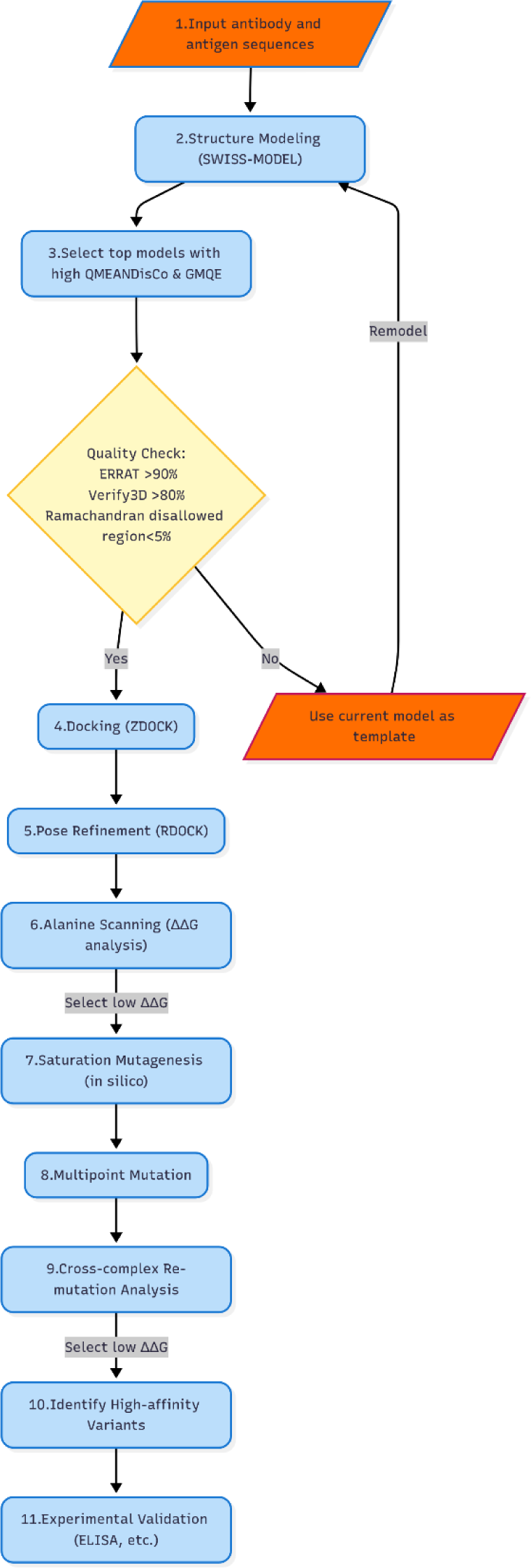
The computational based workflow to enhance binding affinity of antibody.

The quality of the generated models is then assessed using both the GMQE (Global Model Quality Estimation) and QMEAN (Qualitative Model Energy ANalysis) scores. GMQE values range from 0 to 1, with values closer to 1 indicating higher reliability. QMEAN scores typically fall between –4 and 0, where values approaching 0 reflect better agreement between the model and experimentally determined structures. In addition, the QMEANDisCo local plot provides residue-level estimates of model accuracy. Models passing these internal SWISS-MODEL checks are further evaluated using external tools provided by the SAVES server, including ERRAT and Verify3D. Only models with ERRAT scores greater than 90%, Verify3D scores above 80%, and fewer than 5% of residues in disallowed Ramachandran regions are accepted. Models that fail any of these criteria are recycled as templates for a second round of SWISS-MODEL homology modeling, and the quality assessment is repeated until a model meeting all standards is obtained.

Validated antigen and antibody models are then subjected to rigid-body docking using the ZDOCK algorithm. ZDOCK generate a large set of candidate binding poses based on shape complementarity and electrostatic interactions. The top-ranked docking poses are subsequently refined with the RDOCK protocol, which performs energy minimization using the CHARMm force field to optimize van der Waals contacts and electrostatic interactions while estimating desolvation energies. This refinement step filters and re-scores the ZDOCK predictions, increasing the likelihood of identifying near-native antigen–antibody complexes. The final output of these steps is a small number of high-confidence docking conformations that represent plausible binding modes for the nanobody–B7-H3 interaction. These models provide the structural basis for downstream in-silico mutagenesis and affinity-maturation analyses.

In the sixth step, alanine scanning is performed on the antibody–antigen complex to identify potential hotspot residues on the antibody that are critical for binding. This computational approach mutates each residue to alanine and evaluates the resulting change in binding free energy (ΔΔG). Residues that exhibit a significant increase in ΔΔG upon alanine substitution are considered binding “hotspots” as their side chains make key energetic contributions to the interaction. In the following step 7, these identified hotspots undergo in silico saturation mutagenesis. Each hotspot residue is computationally substituted with all other types of amino acids, and the corresponding changes in binding free energy (ΔΔG) are calculated for each variant. The double-point mutation is applied in step 8 to increase the affinity through Synergy. By integrating and comparing the ΔΔG values across multiple predicted complex conformations(poses), the average ΔΔG is used as a robust metric to rank and select the most promising mutations in step 9. The final outcome of this process is a set of single or combinatorial mutations predicted to enhance antibody–antigen affinity in silico, providing rational candidates for experimental validation.

### Antigen-nanobody structural modelling

The antigen B7-H3 sequence and its domain organization are shown in the accompanying figure. Modeling began with SWISS-Model, using templates identified by the built-in homology search. The best initial hit was A0A811YB43.1.A, a hypothetical protein from raccoon dog (*Nyctereutes procyonoides*), which corresponds to the CD276 protein of that species. Notably, this template is not an experimentally determined structure but an AlphaFold v2 prediction. Using this template, an initial model of the antigen (Antigen-1) was generated. However, quality assessment revealed limitations: the template showed an ERRAT score of 91.56 with several residues near position 340 exhibiting high error rates, and only 75.81% of residues achieved a Verify3D score ≥0.1. These values failed to meet the predefined quality thresholds and indicated insufficient structural reliability.

To generate a good model, murine B7-H3 was next selected as a template for SWISS-Model homology modeling. Because this template shares significant homology with both the R1 and R2 domains of human B7-H3, two separate models were built to represent these regions individually. Nevertheless, neither of these domain-specific models passed the quality checks, suggesting that template-based modeling alone was inadequate.

To overcome these limitations, the full-length human B7-H3 structure was predicted directly using AlphaFold, and this AlphaFold model was subsequently used as a template in SWISS-Model for refinement. The refined model achieved an ERRAT score of 91.9 and a Verify3D score above 80%, meeting the quality control criteria. However, the QMEANDisCo Global evaluation indicated poor scores in residues 204–220, a flexible linker connecting the R1 and R2 domains. Such linkers are inherently difficult to predict with high confidence. Considering the prediction challenges in this region, the high sequence homology of R1 and R2, the final working model focused on the R2 region extracted from the refined full-length structure.

**Table 1:**
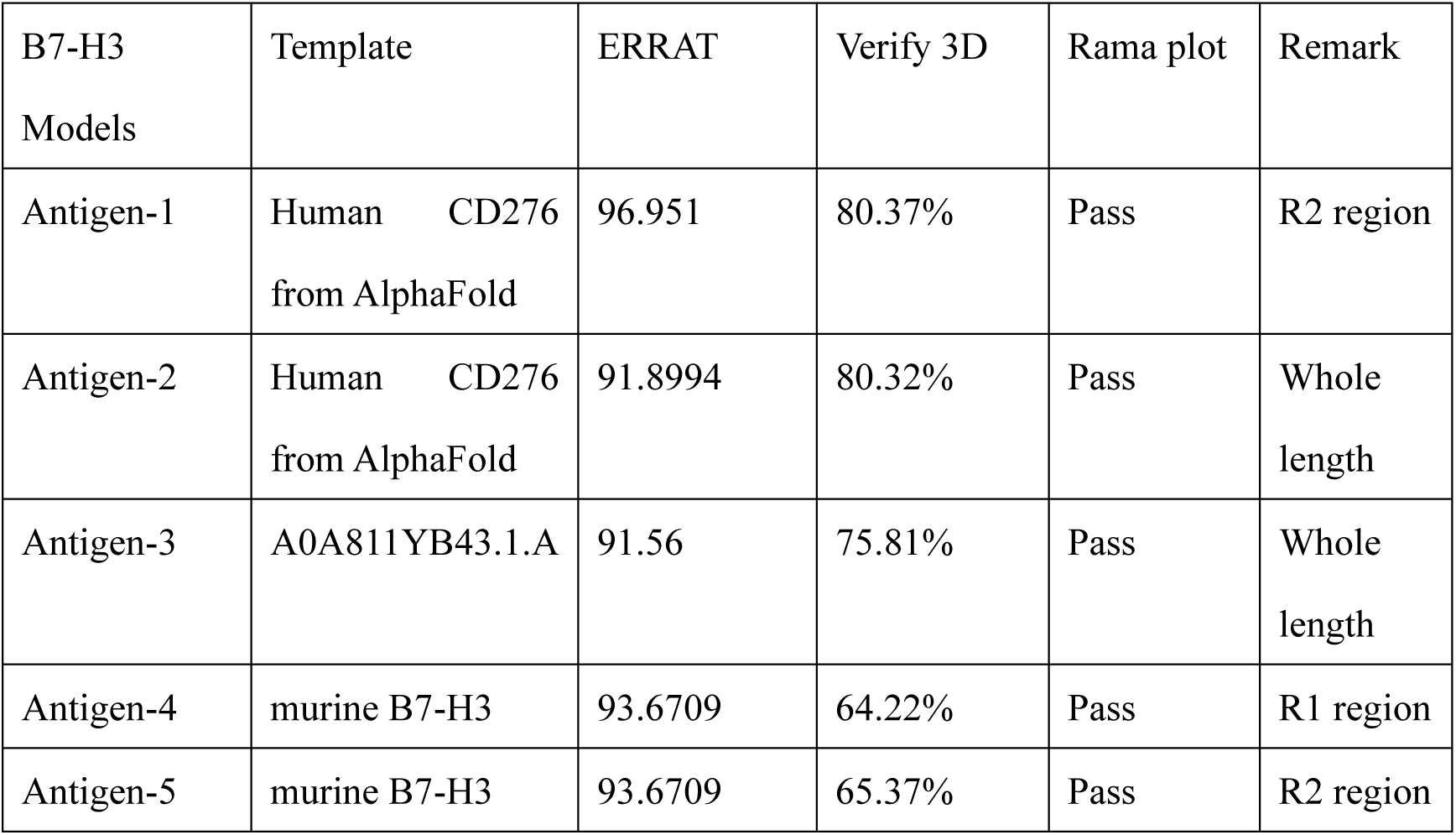
The list of B7-H3 Model candidates. Antigen-1 is selected as final model of B7-H3.

**Figure 2:**
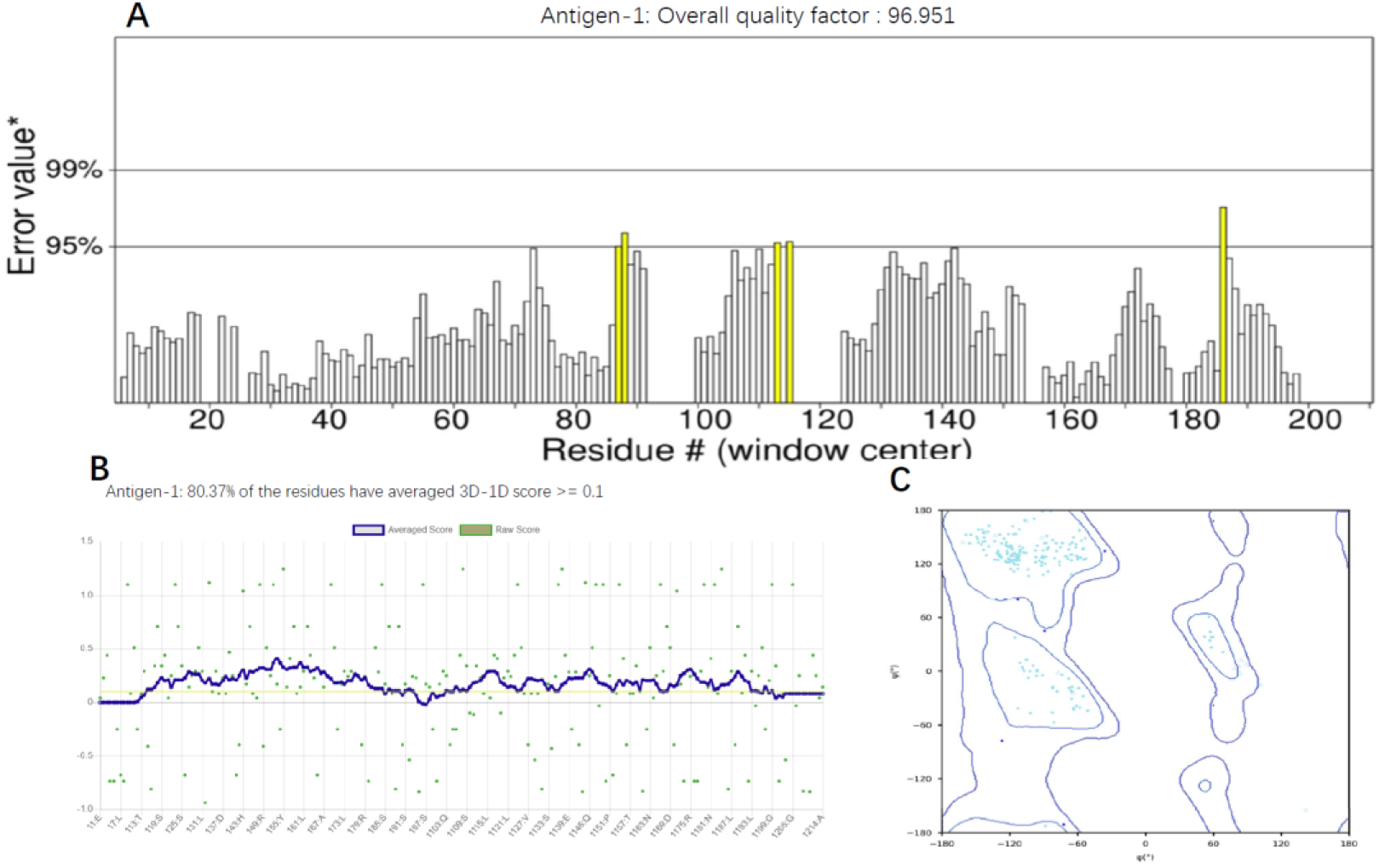
The assessment of Antigen-1(B7-H3) Model. (A) ERRAT overall quality factor of the predicted structure, indicating the non-bonded atomic interaction reliability. (B) VERIFY 3D result showing the percentage of residues with acceptable 3D–1D profiles. (C) Ramachandran plot of the Antigen-1 model, displaying the distribution of residues in favored, allowed, and disallowed regions.

The annotation of the antibody CDR regions was carried out using a hybrid scheme that combines the IMGT, Kabat, and Chothia numbering systems. This approach was chosen to balance the strengths of each scheme: IMGT for its immunogenetics rigor, Kabat for its historical prevalence in antibody studies, and Chothia for its structural relevance. By integrating these systems, the CDR definitions were standardized in a way that ensures both sequence-based and structure-based consistency. Subsequently, homology modeling was performed for the humanized nanobody G1E1. For G1E1, multiple models were generated using SWISS-Model, and their quality was evaluated based on ERRAT, Verify3D, and GMQE scores, as well as structural consistency in the CDR regions. Among the candidates, *model_05* built from the 6m3b.1.D template was selected as the final model. This choice was supported by an ERRAT score of 98, a Verify3D pass rate of 94.4%, and overall favorable quality metrics compared to the other models. In contrast, alternative models such as those based on A0A5C2GG35.1.A, 5csz.1.A, and 8tft templates exhibited localized errors in the CDR2 and CDR3 loops, even when their global scores were reasonable (e.g., GMQE 0.95 for model_01). Thus, while several candidates passed the quality control thresholds, the 6m3b-based model showed the best balance between global quality and local structural accuracy, justifying its selection.

**Figure 3:**
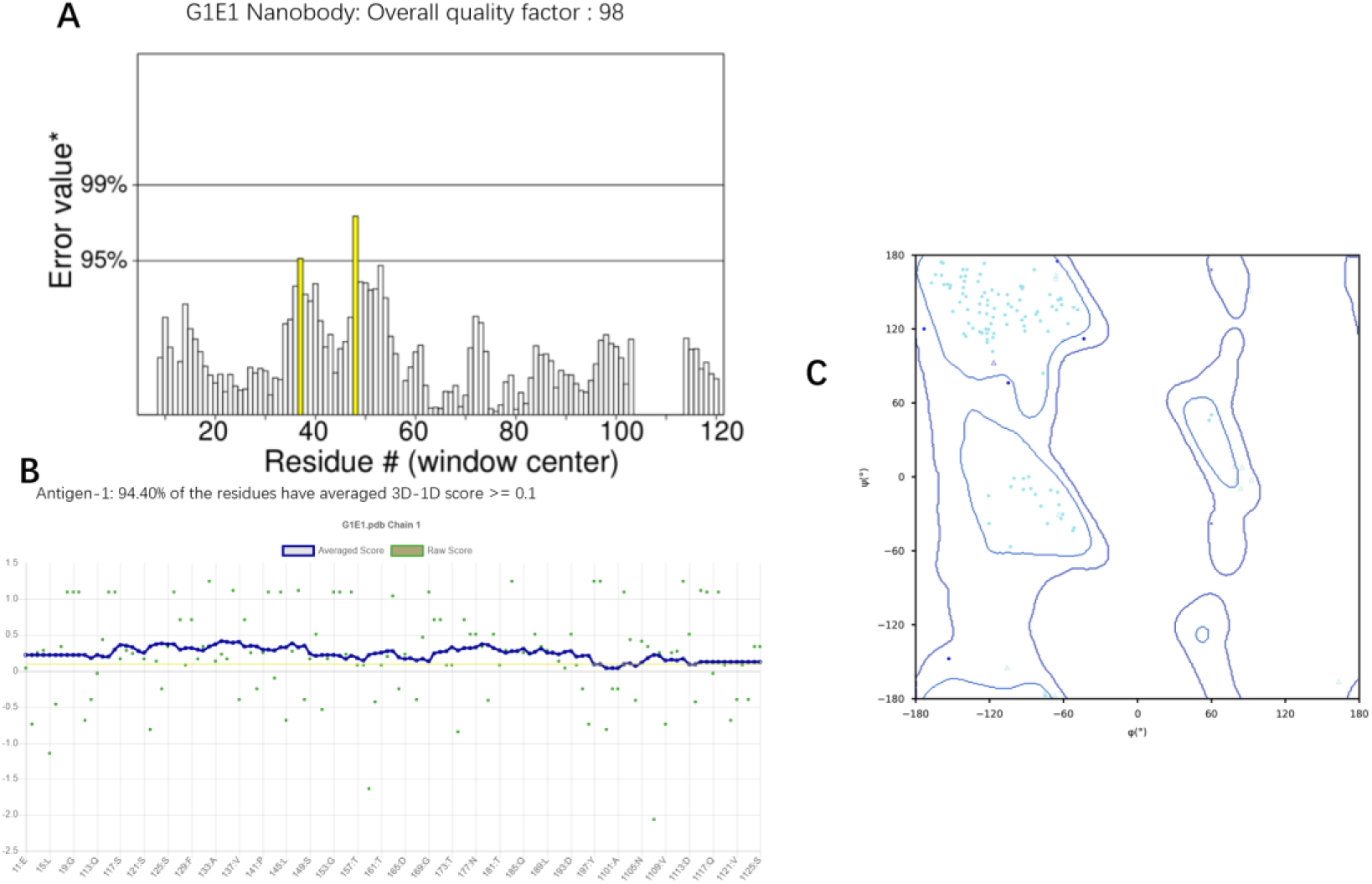
The assessment of model_5 of G1E1 nanobody. (A) ERRAT overall quality factor of the predicted structure, indicating the non-bonded atomic interaction reliability. (B) VERIFY 3D result showing the percentage of residues with acceptable 3D–1D profiles. (C) Ramachandran plot of the Antigen-1 model, displaying the distribution of residues in favored, allowed, and disallowed regions.

Together, these modeling efforts established reliable structural frameworks for both B7-H3 and G1E1, with antigen-1(B7-H3) and model_05 (G1E1)selected as the working models for downstream docking and mutagenesis studies.

### Docking of B7-H3 and G1E1, B4-6 Nanobodies

ZDock was employed to perform the docking between the nanobody models and the B7-H3 antigen. The program provides flexible parameter options, including the ability to define residues on the receptor or ligand that should either participate in binding or be blocked from interaction. In this study, the nanobody model G1E1 were used as receptors, while the B7-H3 R2 model was designated as the ligand.

Based on prior flow cytometry results(Figure 4), it was known that the G1E1 nanobody predominantly recognizes the C2 subregion within the R2 domain. To capture both unbiased and targeted docking behaviors, two types of docking were carried out: one with no spatial constraints and another with restraints enforcing binding within the experimentally suggested epitope regions. As a result, 2 initial complex datasets were generated: *ZDock-G1E1-B7H3R2C2*, and *ZDock-G1E1-B7H3R2 (unrestricted)*. For each docking run, ZDock generated 2,000 candidate poses(Figure 5).

**Figure 4:**
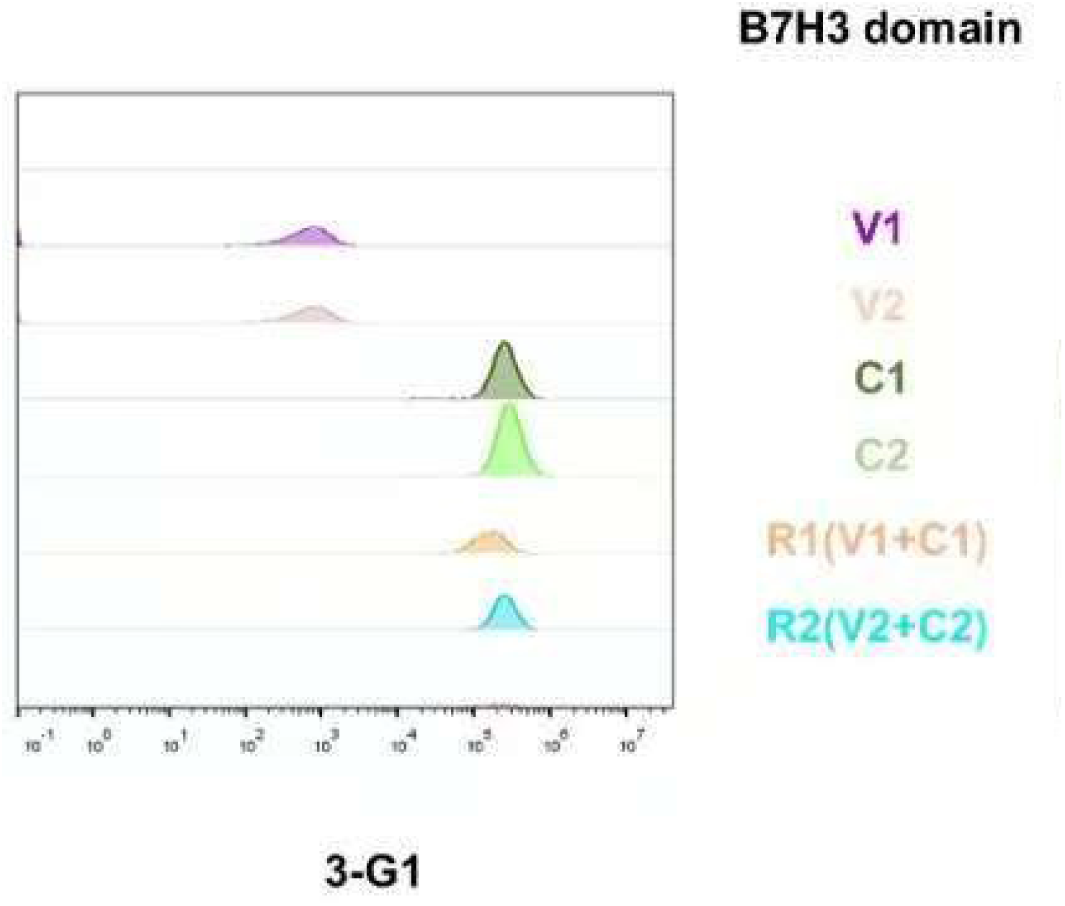
The flow cytometry results of G1-cam antibody. It can be seen that the antibody mainly binds to the C regions of the antigen.

**Figure 5:**
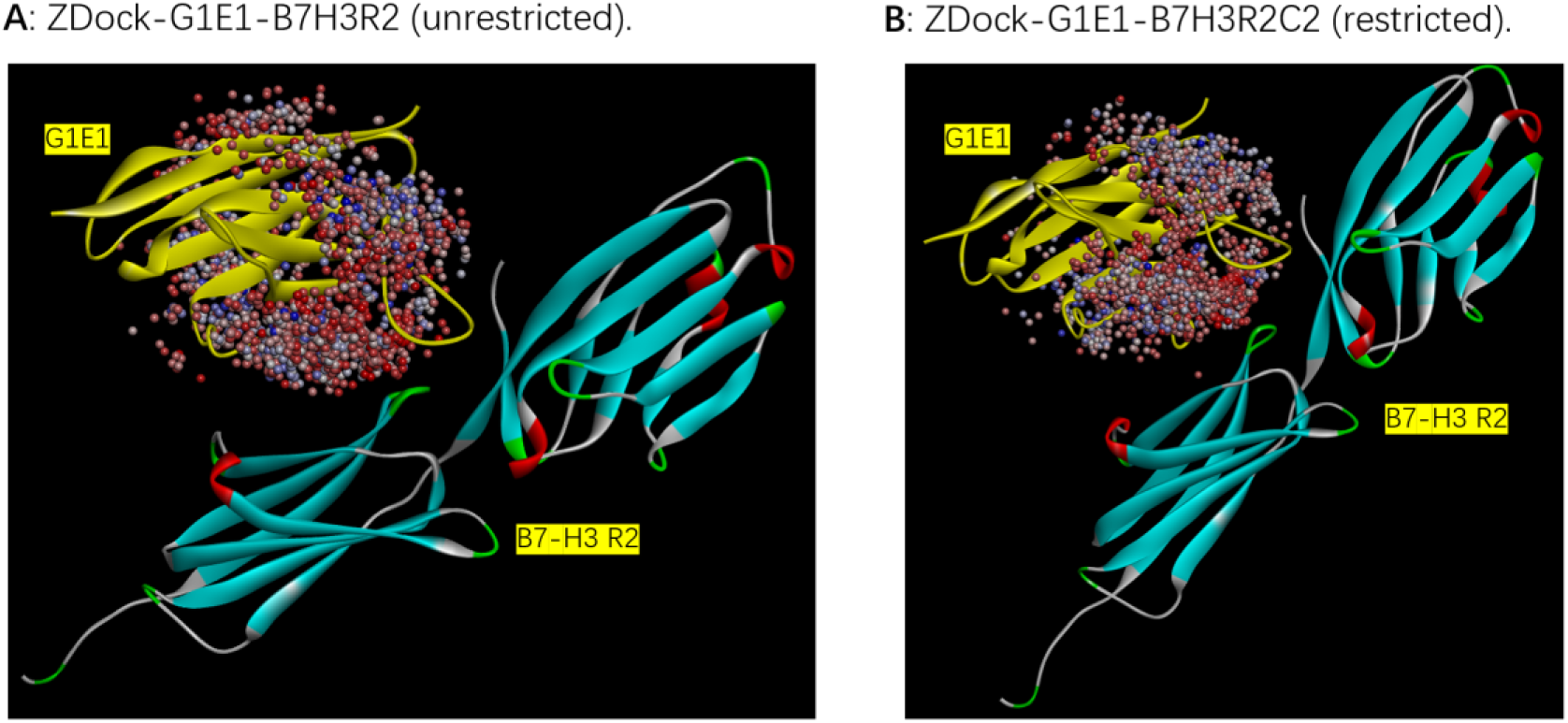
Output of ZDock Process. The yellow structure represents the G1E1 nanobody, while the other chain corresponds to the R2 region of B7-H3. Each dot indicates one predicted docking pose of the nanobody, and a total of 2000 poses were generated by ZDOCK.

The 2,000 raw poses in 200 representative clusters from each docking run were subsequently selected through *Top poses in Largest Cluster* and refined using the process_poses function within ZDock(Figure 6). Finally, the resulting complexes were further refined using the RDOCK program, which applies energy minimization and scoring functions to discriminate near-native complexes from spurious ones. The RDOCK step produced high-confidence antigen–nanobody complex structures that could then be used as the starting point for in silico mutagenesis and affinity maturation studies.

**Figure 6:**
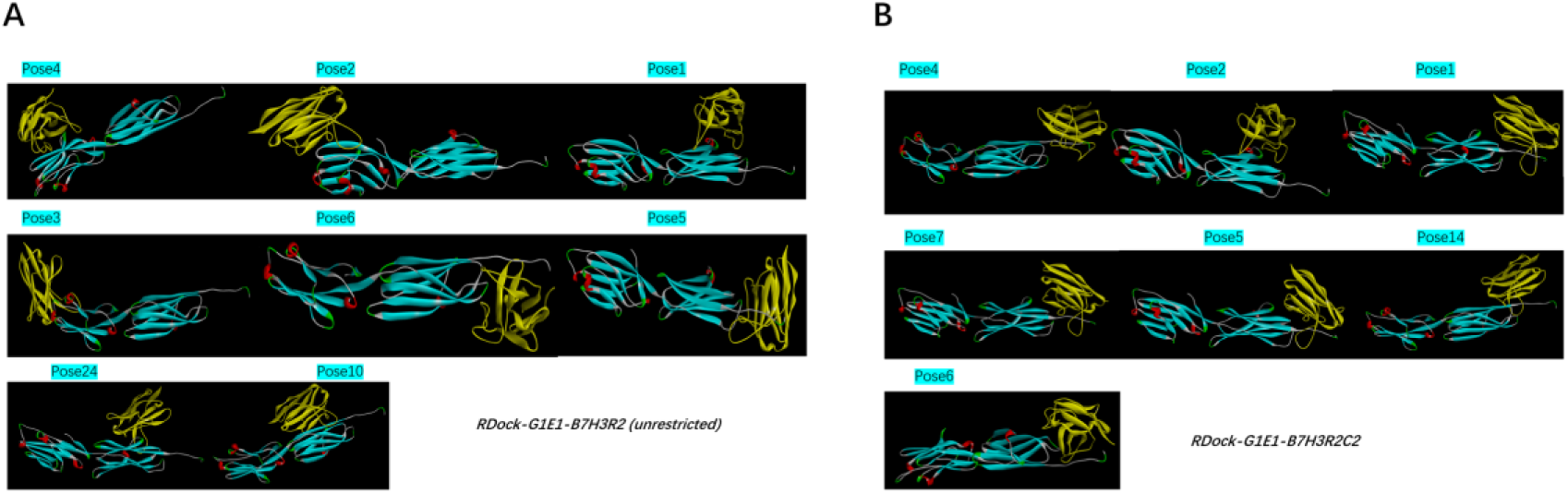
The outputs of Rdock process. The yellow structure represents the G1E1 nanobody, while the other chain corresponds to the R2 region of B7-H3. (A). 8 poses selected for G1E1-B7H3R2 complex(unrestricted). (B). 7 poses for G1E1-B7H3R2 complex(C2 restricted)

### Alanine scanning and Mutation in silico

Alanine scanning was first applied to the antigen–nanobody complexes to identify potential hotspot residues within the nanobody that contribute significantly to binding affinity. This computational technique systematically mutates each residue of Antibody to alanine and calculates the resulting change in binding free energy (ΔΔG). Residues exhibiting the most negative ΔΔG values were considered key contributors to antigen recognition. The results were ranked from low to high ΔΔG, and the top ten residues were selected as potential mutational hotspots for subsequent analysis. The output is listed in Supplement 1.

Following alanine scanning, saturation mutagenesis was performed on each of these hotspot residues by substituting them with all 20 standard amino acids. The resulting mutants were again evaluated based on ΔΔG values, with mutations yielding ΔΔG values less than −1 kcal/mol considered likely to enhance binding affinity. Artificial residues such as ASPH or noncanonical modifications provided by the modeling software were excluded from consideration during data processing. Every pose from each of the RDOCK-refined complexes underwent this two-step mutation analysis, and the results were summarized and sorted according to the predicted affinity improvement in supplement 1. The promising mutations identified in this step were then subjected to stability prediction to ensure that enhanced binding affinity would not compromise the nanobody’s structural integrity. Mutations predicted to reduce overall thermal stability or those located outside the complementarity-determining regions (CDRs) were excluded. This exclusion criterion is not because framework residues are functionally irrelevant, but rather due to the higher risk that framework alterations could disrupt the nanobody’s overall fold and solubility, while CDR mutations typically preserve structural integrity.

To further optimize binding, selected single-point mutations were analyzed in the Discovery Studio (DS) mutation module to evaluate the effects of double-point mutations. In this step, DS automatically generated all possible pairwise combinations of the selected single-site mutations and calculated their predicted impact on binding free energy. Although multiple simultaneous mutations could theoretically yield higher affinity, complex residue interactions and the inherent limitations of in silico prediction models could lead to structural artifacts. Therefore, the double-point mutation approach was chosen as a balanced strategy that combines the potential for meaningful affinity enhancement with manageable structural perturbation. The ΔΔG results of all double mutants were ranked in ascending order, and the top candidates for both nanobody–antigen complexes were listed in the supplement 2.

### Re-mutation Analysis

In the re-mutation analysis, a comprehensive reassessment was performed based on the dataset generated from the previous step, which contained all mutation combinations predicted to enhance binding affinity across different docking poses. These poses were selected from the largest docking clusters or from high-scoring, less-populated clusters(cluster 2001), each representing a distinct potential binding conformation between the nanobody and the antigen. By examining mutations that consistently improved affinity across multiple poses, we identified a subset of mutations that demonstrated robust affinity enhancement regardless of docking conformation.

To further refine the mutation screening, all poses were subjected to a second round of double-point mutation analysis. This step was necessary because some residues were not included in the initial alanine scanning results for certain poses. Alanine scanning serves as a rapid and computationally efficient method to identify potential mutation hotspots, but it inherently limits the search space. Performing full saturation mutagenesis on every residue would provide the most comprehensive results but is computationally impractical. As a result, alanine scanning may overlook residues that show little change upon alanine substitution yet could yield substantial affinity improvements when mutated to other amino acids. Conversely, residues identified as hotspots by alanine scanning may not necessarily produce favorable effects in all cases. To address these limitations, we integrated data across multiple poses and re-applied selected mutation combinations to all of them. This cross-pose approach helped reduce bias introduced by pose-specific differences and compensated for omissions from the alanine scan. By comparing ΔΔG variations across poses, we were able to cross-validate the robustness of each mutation’s predicted affinity improvement and identify mutations with consistent favorable effects.

The ΔΔG values from the re-mutation simulations were collected and averaged for each mutation combination. For docking clusters containing multiple poses, the mean ΔΔG of those poses was taken as the representative value for that cluster. In contrast, for most smaller clusters containing only a single pose, that individual ΔΔG value was directly used. It is worth noting that cluster 2001 represented discrete, high-scoring poses that did not belong to any of the main clusters but were retained due to their strong docking scores and structural plausibility. Finally, mutation combinations exhibiting the lowest average ΔΔG values—indicating the greatest predicted affinity improvements—were shortlisted. Two double-point mutations showing the most consistent and significant affinity enhancement across poses were selected as the final candidates for subsequent experimental validation. The result of re-mutation is listed in supplement 3.

The most abundant band corresponds to the 16 kDa nanobody; however, several additional minor protein bands were also observed in the samples. Since these extra bands appeared consistently across all four samples, they are likely endogenous *E. coli* proteins derived from the expression system. These bacterial proteins may possess intrinsic affinity for Ni²⁺ ions, leading to their co-purification with the nanobody during Ni–NTA affinity chromatography.

### SEC Analysis and further purification

Size-exclusion chromatography (SEC) was employed to assess the purity and homogeneity of the proteins following Ni–NTA affinity purification. The chromatograms of all four samples exhibited a dominant peak corresponding to the nanobody. In addition to the major peak, several minor peaks with lower mAU(MilliAbsorbance Unit) were detected, indicating the presence of trace impurities. Interestingly, the elution patterns of these minor peaks were similar across all samples, proving the previous guess that the impurities are not specific to individual constructs but rather represent host-derived E. coli proteins that co-purified. To further evaluate the purification efficiency, fractions from the main SEC peak(Figure 7) and the unprocessed samples were analyzed by Native-PAGE(Figure 8). The nanobody band was markedly enriched after SEC, while the intensity of non-specific bands was substantially reduced, confirming that SEC effectively removed residual contaminants and yielded a highly purified nanobody preparation suitable for downstream analyses.

**Figure 7:**
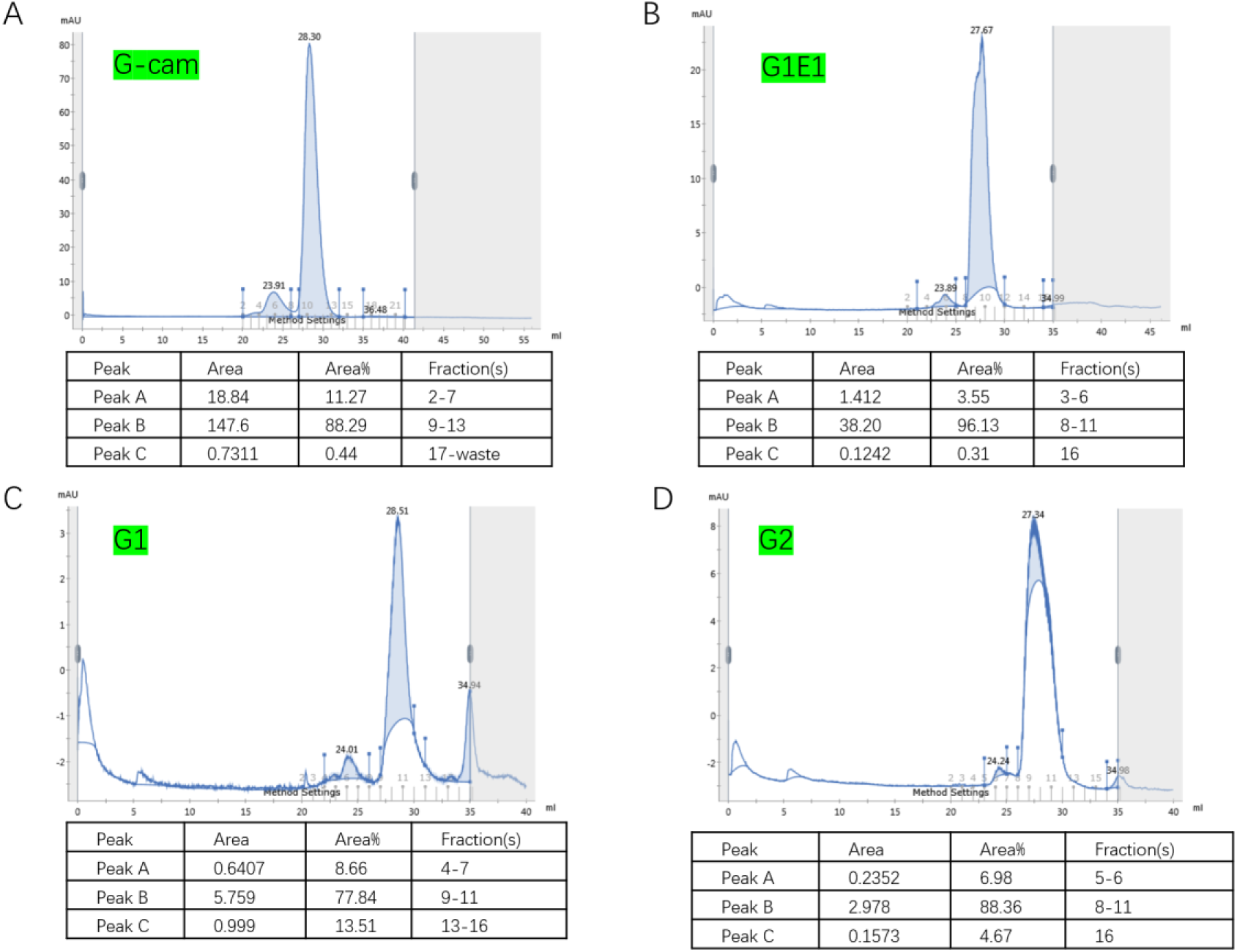
The result of size-exclusion chromatography (SEC). The X-axis represents the elution volume, and the Y-axis represents the absorbance (mAU). The major peak corresponds to the nanobody, and its area percentage reflects the relative abundance and purity of the target protein.

**Figure 8:**
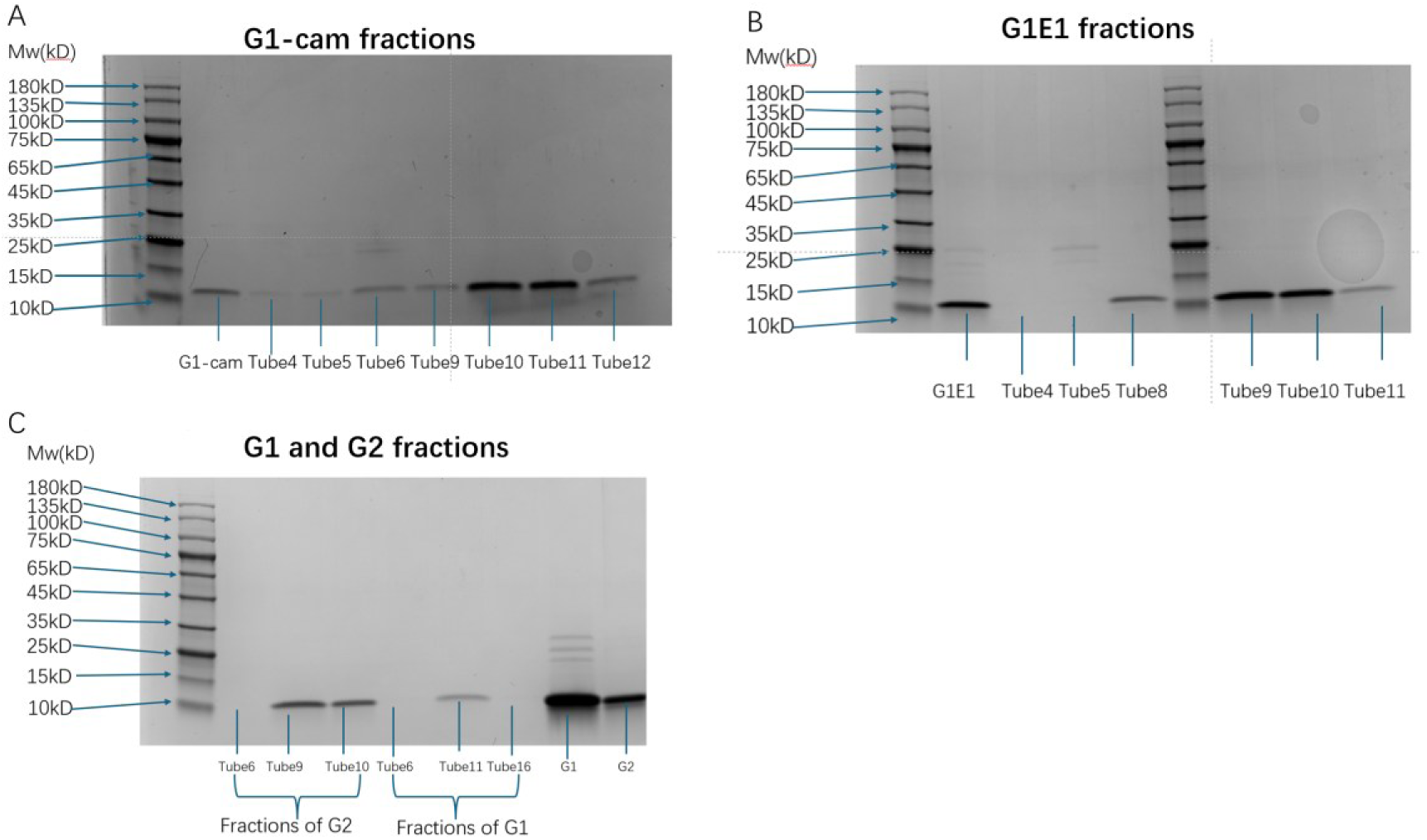
Native-PAGE analysis of SEC fractions for four nanobodies. Each lane corresponds to one SEC fraction (Tube). Several fractions were excluded from analysis because Nanodrop measurements indicated negative or negligible protein concentrations. (A) Native-PAGE results for G1-cam and its SEC fractions. From left to right: molecular weight marker, unpurified G1-cam, Tube 4, Tube 5, Tube 6, Tube 9, Tube 10, Tube 11, and Tube 12. (B) Native-PAGE results for G1E1 and its SEC fractions. From left to right: molecular weight marker, unpurified G1E1, Tube 4, Tube 5, Tube 8, molecular weight marker, Tube 9, Tube 10, and Tube 11. (C) Native-PAGE results for G1 and G2 nanobodies. From left to right: molecular weight marker, G2 fractions (Tube 6, Tube 9, Tube 10), G1 fractions (Tube 6, Tube 11, Tube 16), unpurified G1, and unpurified G2.

### Validation of affinity by ELISA

In the ELISA assay, G1E1 displayed a markedly reduced binding affinity to B7-H3 compared with the original G1-cam, reflecting the affinity loss commonly associated with antibody humanization. In contrast, both engineered variants, G1 and G2, exhibited enhanced binding affinity, with G1 showing the most significant improvement. The corresponding ELISA binding curves and estimated EC50 values are presented below to illustrate these differences quantitatively(Figure 9).

**Figure 9:**
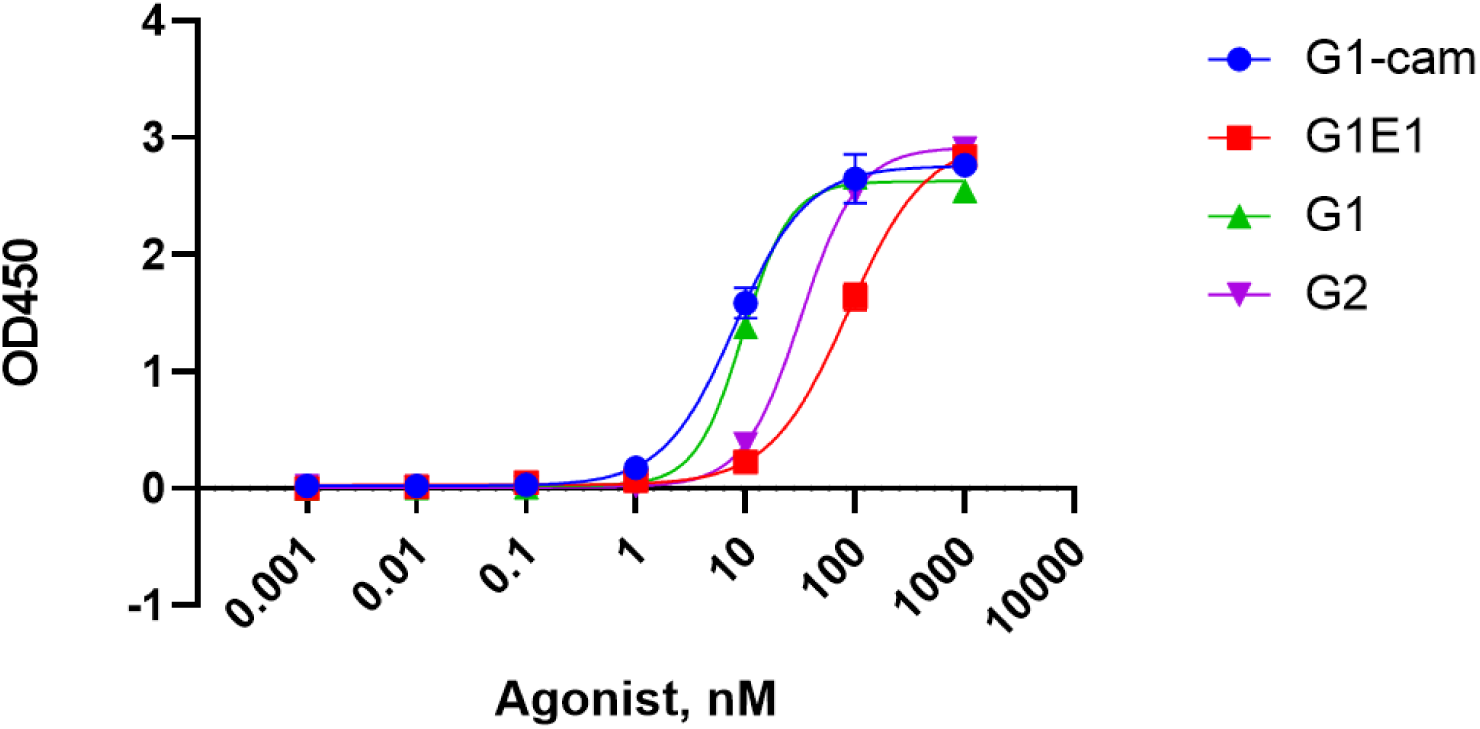
The ELISA result of four antibodies and the antigen is B7-H3. And EC50(nM) of each nanobody is showed in table.

### Biolayer Interferometry (BLI) Analysis

The binding kinetics of the nanobody G1 and its humanized variant G1E1 toward the target antigen B7-H3 were evaluated using biolayer interferometry (BLI). Both antibodies exhibited typical association and dissociation profiles, indicating specific binding to the antigen. The kinetic parameters obtained from the 1:2 binding model are summarized in Table 3.

**Figure 10:**
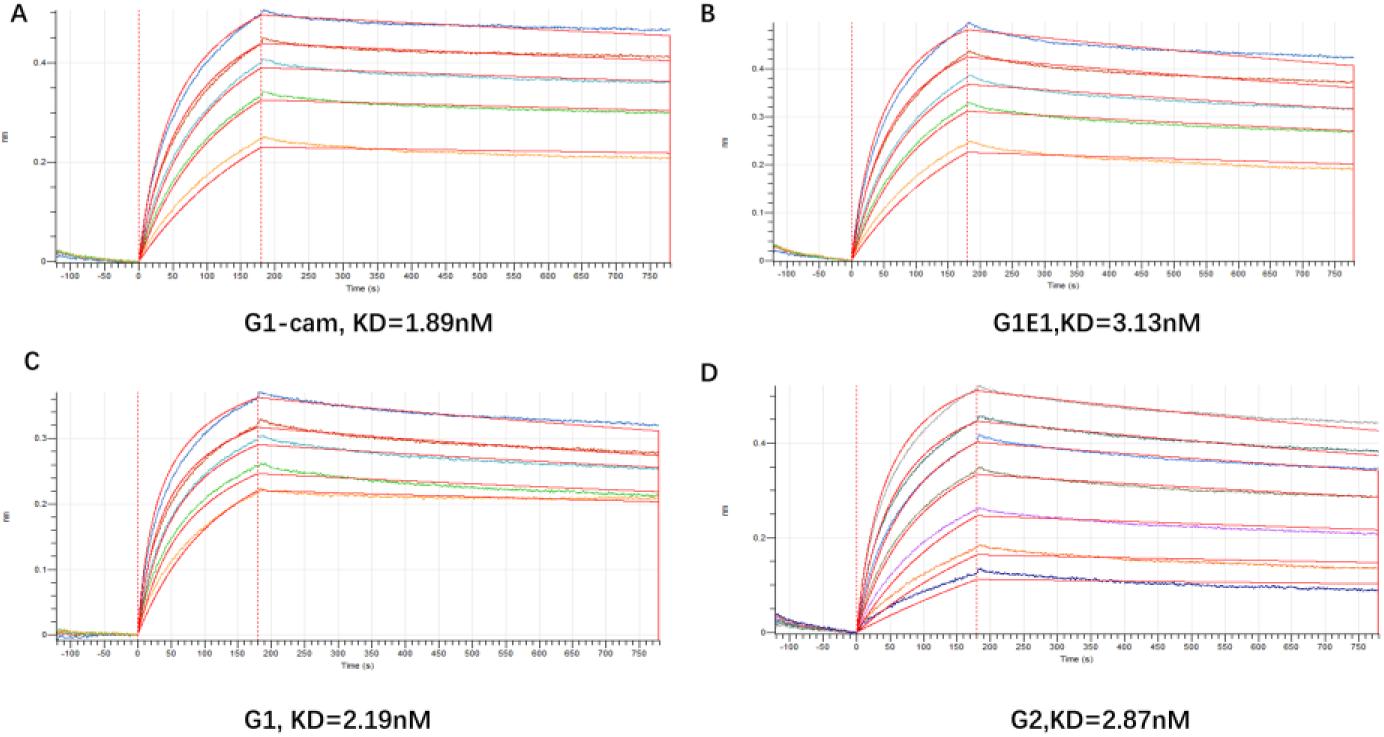
The association and dissociation curves obtained from BLI analysis. The experiment consisted of three sequential phases: a 120-second baseline phase, followed by a 180-second association phase, and a 600-second dissociation phase. The sensorgrams illustrate the real-time binding and unbinding kinetics of the nanobody–antigen interactions.

**Table 2:** This table shows the ddG and Mutations of 2 selected variants G1 and G2 under 2 different docking restrictions.

| RDock | Combination | Mean ddG | Name |
| --- | --- | --- | --- |
| **G1E1-B7H3R2C2** | ASP65ARG+GLU111LYS | -3.037 | G1 |
|  | VAL109ARG+GLU111ARG | -2.93 | G2 |
| **G1E1-B7H3R2** | ASP65ARG+GLU111LYS | -3.22429 | G1 |
|  | VAL109ARG+GLU111ARG | -3.22071 | G2 |

**Table 3:** Kinetic parameters were derived from global fitting of the BLI sensorgrams.

| Antibodies | $K_a(1/Ms) \times 10^4$ | $K_{dis}(1/s) \times 10^{-4}$ | $K_D(M) \times 10^{-9}$ |
| --- | --- | --- | --- |
| **G1-cam** | 9.3870 | 1.777 | 1.8930 |
| **G1E1** | 10.55 | 3.352 | 3.1777 |
| **G1** | 15.03 | 3.289 | 2.1880 |
| **G2** | 12.18 | 3.5 | 2.8740 |

The equilibrium dissociation constant (Kᴅ) values revealed a measurable improvement in affinity for the designed variants compared to the humanized antibody G1E1. Specifically, the G1 variant exhibited a Kᴅ of 2.19 × 10⁻⁹ M, representing an approximately 1.45-fold the binding affinity of G1E1 (3.18 × 10⁻⁹ M). Similarly, the G2 variant showed a modest improvement, with a Kᴅ of 2.87 × 10⁻⁹ M, corresponding to 1.11-fold affinity of G1E1. In terms of kinetic parameters, both G1 and G2 displayed higher association rate constants (Kₐ) compared with G1E1, suggesting faster antigen recognition. The dissociation rate constants (Kdis) remained closed among all variants, indicating that the mutations primarily enhanced the association process rather than reducing the dissociation rate. Collectively, these results demonstrate that the in silico-guided mutations improved binding kinetics, yielding antibodies with stronger and faster antigen engagement while maintaining overall binding stability.

### Thermal Stability Analysis by Circular Dichroism (CD)

The thermal stability of the antibodies was evaluated using circular dichroism (CD) spectroscopy. Thermal unfolding curves were recorded to determine the melting temperature (Tₘ) and van’t Hoff enthalpy (ΔHᵥᴴ) of each protein. As summarized in Table 4, the humanized antibody G1E1 exhibited a melting temperature of 56.3991 °C, while the parental antibody G1 displayed a Tₘ of 56.5619 °C.

**Table 4:** The melting temperature (Tₘ) and van’t Hoff enthalpy (ΔHᵥᴴ) of antibodies.

| Antibody | $\Delta H_v^H$ | $T_m(^{\circ}\text{C})$ |
| --- | --- | --- |
| G1E1 | 128.073 | 56.3991 |
| G1 | 131.795 | 56.5619 |

A slight increase in thermal stability was observed following affinity-enhancing mutation, the difference between the two proteins was relatively small. This indicates that the in silico–guided mutagenesis preserved the overall structural integrity of the antibody, ensuring that the improved affinity did not come at the cost of substantial destabilization. These findings suggest that the computationally predicted substitutions are both functionally effective and structurally tolerable, supporting the robustness of our design workflow.

## Discussion

In this study, we established a standardized and reproducible *in silico* protocol for antibody affinity maturation using a B7-H3–nanobody system as example. The workflow integrates molecular docking, alanine scanning, saturation mutagenesis, and ΔΔG-based affinity evaluation to identify mutations that enhance antigen binding while maintaining structural stability. By systematically combining multiple docking poses, cross-validating ΔΔG results across conformational ensembles, and filtering mutations through thermodynamic and structural criteria, this pipeline provides a rational and efficient strategy to predict affinity-enhancing mutations prior to experimental validation.

A key advantage of this protocol is its balance between computational feasibility and biological relevance. Instead of performing exhaustive saturation mutagenesis on every residue—which would be computationally prohibitive—we employed alanine scanning as a pre-filter to rapidly locate potential hotspots. Subsequent targeted saturation mutagenesis around these positions, followed by multi-pose re-mutation analysis, helped recover beneficial mutations that might otherwise be missed by conventional hotspot-based approaches. This layered strategy not only improved the robustness of the prediction but also reduced the bias introduced by single-pose or interface-restricted analyses. The final mutations identified by this protocol demonstrated experimentally measurable improvements in binding affinity, validating the predictive power and general applicability of the method.

Notably, ELISA and BLI assays exhibited partially divergent results regarding the magnitude of affinity improvement. While ELISA measurements showed a substantial enhancement in apparent binding potency, with the EC₅₀ decreasing from 86.12 nM to 9.545 nM(G1 nanobody) and 86.12nM to 32.02nM(G2 nanobody), the kinetic analysis by BLI revealed a strong change in equilibrium dissociation constant (Kᴅ) from 3.13 nM to 2.19 nM and a modest change in Kᴅ, from 3.13nM to 2.87nM. This discrepancy can be attributed to the inherent methodological differences between the two assays. ELISA reflects an endpoint measurement influenced by avidity effects, surface immobilization, and potential conformational stabilization of the antibody–antigen complex on a solid phase, which can amplify apparent binding differences.

In contrast, BLI measures real-time kinetic interactions in solution under near-equilibrium conditions, providing a more direct quantification of intrinsic affinity. Therefore, the pronounced difference observed in ELISA likely reflects enhanced binding affinity or slower dissociation in a surface-bound context, rather than a dramatic change in the intrinsic affinity constant. Such context-dependent effects highlight the importance of interpreting ELISA-derived potency as a composite of affinity, avidity, and particularly for nanobody–antigen interactions.

The thermal stability analysis further supports the structural integrity of the engineered antibodies. Circular dichroism spectroscopy revealed that the melting temperature (Tₘ) of the optimized variant showed only a minor increase compared with the parental antibody. This minimal change indicates that the introduced mutations did not substantially disturb structural stability of the nanobody framework while enhancing binding affinity. Therefore, our in silico– guided design effectively enhanced antigen affinity while preserving the conformational robustness of the antibody.

As previously mentioned, B7-H3 was deliberately chosen as a particularly challenging model antigen due to the absence of any experimentally determined crystal structure. The successful application of our computational workflow under such structurally limited conditions highlights its robustness and adaptability. Despite the inherent uncertainty of homology-based modeling, the predicted mutations still led to measurable improvements in binding affinity, demonstrating that the workflow remains reliable even without high-resolution structural information. This outcome underscores the practical value of our in silico protocol for antibody engineering against structurally unresolved or difficult-to-model targets. Beyond the specific case of anti-B7-H3 nanobodies, the established protocol offers a versatile framework for computational antibody engineering. The modular design allows for easy adaptation to different antibody formats or target antigens, including those of therapeutic importance such as immune checkpoints and tumor-associated molecules. Given that B7-H3 has been implicated in tumor immune evasion, metastasis, and therapy resistance, improving the affinity and specificity of anti-B7-H3 antibodies holds significant translational potential for cancer immunotherapy. Moreover, the computational standardization achieved here paves the way for large-scale *in silico* screening and rational antibody design, reducing experimental cost and time while maintaining predictive accuracy.

In conclusion, this work not only provides insight into the structural determinants of B7-H3 recognition but also establishes a generalized and standardized *in silico* workflow for antibody affinity optimization. Future work will focus on integrating molecular dynamics simulations and machine-learning-based scoring models to further refine mutation predictions and extend this framework to other clinically relevant antigen.

